# Screening of SARS-CoV-2 Antivirals Through a Cell-Based RNA-Dependent RNA Polymerase (RdRp) Reporter Assay

**DOI:** 10.1101/2022.04.04.486994

**Authors:** Timsy Uppal, Kai Tuffo, Svetlana Khaiboullina, Sivani Reganti, Mark Pandori, Subhash C. Verma

## Abstract

COVID-19 (Coronavirus Disease 2019) caused by SARS-CoV-2 (Severe Acute Respiratory Syndrome CoronaVirus-2) continues to pose international public health threat and thus far, has resulted in greater than 5.6 million deaths worldwide. Vaccines are critical tools to limit COVID-19 spread, but antiviral drug development is an ongoing global priority due to fast spreading COVID-19 variants that may elude vaccines efficacies. The RNA-dependent RNA polymerase (RdRp) of SARS-CoV-2 is an essential enzyme of viral replication and transcription machinery complex. Therefore, the RdRp is an attractive target for the development of effective anti-COVID-19 therapeutics. In this study, we developed a cell-based assay to determine the enzymatic activity of SARS-CoV-2 RdRp through luciferase reporter system. The SARS-CoV-2 RdRp reporter assay was validated using a known inhibitors of RdRp polymerase, remdesivir along with other anti-virals including ribavirin, penciclovir, rhoifolin, 5’CT, and dasabuvir. Among these inhibitors, dasabuvir (FDA-approved drug) exhibited promising RdRp inhibitory activity. Anti-viral activity of dasabuvir was also tested on the replication of SARS-CoV-2 through infection of Vero E6 cells. Dasabuvir inhibited the replication of SARS-CoV-2, USA-WA1/2020 as well as B.1.617.2 (delta variant) in Vero E6 cells in a dose-dependent manner with IC_50_ values 9.47 μM and 10.48 μM, for USA-WA1/2020 and B.1.617.2 variants, respectively). Our results suggests that dasabuvir can be further evaluated as a therapeutic drug for COVID-19. In addition, our assays provide robust, target-specific, and high-throughput screening compatible (z- and z’-factors of > 0.5) platforms that will be a valuable tool for the screening SARS-CoV-2 RdRp inhibitors.

**Significance:** SARS-CoV-2 has caused a major public crisis world has seen in recent history. Development of vaccines and emergency use authorization of anti-virals are helping in reducing the burden of SARS-CoV-2 caused hospitalization and deaths. However, there is still need for optimal anti-viral(s) that can efficiently block viral propagation, and targeting viral polymerase (RdRp) is an among the most suitable targets for clamping viral replication. In this study, we developed a cell-based assay to screen potential compounds capable of blocking RdRp activity. The efficacy of our assay was validated by using already approved anti-virals, which reduced RdRp activity and slowed the replication of two SARS-CoV-2 variants (WA1 USA-WA1/2020 and B.1.617.2) in a cell culture model. This confirmed that our system can be used for identifying potential anti-SARS-CoV-2 anti-virals.

## 1. Introduction

The life-threatening COVID-19 is caused by SARS-CoV-2, a new coronavirus discovered in 2019, which caused a global pandemic (1). SARS-CoV-2 belongs to the *betacoronavirus* genus, together with SARS-CoV and MERS-CoV, and has high potential to cause lethal zoonotic infections among existing HCoVs (2). SARS-CoV-2 mainly attacks the lower respiratory system to cause pneumonia-like symptoms, but can also damage digestive system, gastrointestinal system, heart, kidney, liver, and central nervous system leading to multiple organ failure and death in severe COVID-19 patients (3). Extensive efforts are being made worldwide to develop efficacious antiviral drugs against COVID-19 infections.

SARS-CoV-2 is an enveloped, positive-sense single-stranded RNA virus. The polycistronic RNA genome (~30kb) has 14 open reading frames (ORFs) that encode for two groups of proteins (4). Approximately, one-third of SARS-CoV-2 genome consists of four structural proteins [spike (**S**), envelope I, membrane (**M**), and nucleocapsid (**N**)] and six accessory proteins, encoded by ORFs 2-10 (5). The remaining two-third of viral genome, comprises of non-structural proteins, expressed as two large replicase polyproteins (ORF1a and ORF1ab) which are cleaved by virus’s proteases, M^pro^ and 3CL^pro^ into 16 small peptides, nsp1-16. These replicase proteins assemble into a membrane-bound ribonucleoprotein complex (or replication and transcription complex) that directs RNA synthesis and processing. Several proteins of nsp interactome thus serve as potential targets for the development of antivirals (6). The RNA-dependent RNA polymerase (RdRp), one of the SARS-CoV-2 non-structural proteins (nsp12), is the central catalytic subunit of the viral RNA synthesizing machinery (7). The crucial role in the virus replication and absence of closely related homolog in humans makes RdRp an important druggable molecular target of SARS-CoV-2 inhibition.

Hillen *et al* recently resolved the structure of replicating SARS-CoV-2 RdRp polymerase in its active form using cryo-electron microscopy (8). Similar to RdRp of SARS, SARS-CoV-2 RdRp also consists of catalytic domain of nsp12, one copy of nsp7, two copies of nsp8, and more than two turns of RNA template-product duplex. As a result, known inhibitors of RdRp of SARS might show promising effect against RdRp of SARS-CoV-2. A wide array of adenine, and guanine-based nucleoside analog inhibitors targeting viral RdRp polymerase have been considered against SARS-CoV-2 (9, 10). However, currently, remdesivir (GS-5734) by Gilead Sciences, molnupiravir by Merck and paxlovid by Pfizer are the only FDA emergency use authorized drugs for the treatment of mild to moderate COVID-19 infection (11). These drugs are shown to inhibit SARS, MERS and SARS-CoV-2 replication in cell culture and animal models by either targeting RdRp polymerase (12–15) or M^pro^ main protease (16). Since virus accumulates mutations following replication, there is a need for the development of anti-virals capable of blocking multiple targets with better potency and selectivity.

Identification of specific inhibitors through high-throughput screening (HTS) requires a cost-efficient, robust, convenient to use, and highly reliable assay. To improve current efforts to find SARS-CoV-2 antivirals, especially RdRp-specific inhibitors, in this study, we developed a HEK293 cell-based stable RdRp reporter assay capable of specifically monitoring SARS-CoV-2 RdRp polymerase activity in cellular conditions. The SARS-CoV-2 RdRp reporter was developed by modifying the reporter systems reported for HCV NS5B, and MERS-CoV (17, 18). The use of internal control monitoring RdRp polymerase activity allowed for the identification of selective RdRp inhibitors in a single assay. We further evaluated the inhibitory effect of known RdRp inhibitors, i.e., rhoifolin, dasabuvir, ribavirin, penciclovir, and cytidine-5’triphosphate on SARS-CoV-2 RdRp activity. Using the optimized cell-based reporter system, we, confirmed dasabuvir as a direct SARS-CoV-2 RdRp inhibitor, demonstrating the efficacy of our RdRp screening assay. Furthermore, these findings were validated using SARS-CoV-2 infected Vero E6 cells. dasabuvir alone as well as in combination with Ribavirin potently inhibited SARS-CoV-2 replication in USA-WA1/2020 infected Vero E6 cells. Thus, our results demonstrated that dasabuvir is a potent SARS-CoV-2 replication inhibitor. Also, the developed SARS-CoV-2 RdRp reporter assay could be used in high-throughput screening to discover compounds specific to SARS-CoV-2 RdRp with low cytotoxicity and reduced side-effects.

## 2. Materials and Methods

### 2.1. Cells, Reagents and Antibodies

The human embryonic kidney HEK293 and Vero E6 cells were obtained from ATCC and maintained in Dulbecco’s modified Eagle medium (DMEM) supplemented with 10% FBS (Atlanta Biologicals), 2 mM L-glutamine, 25 U/mL penicillin, and 25 μg/mL streptomycin. The cells were grown at 37 °C and 5% CO2 in a humidified chamber. G418 and Hygromycin were purchased from InvivoGen (San Diego, CA, United States). Inhibitors, namely remdesivir (GS-5734), rhoifolin, ribavirin, dasabuvir, penciclovir, and cytidine-5’-triphosphate were purchased from Selleck Chemicals LLC (Houston, TX, United States) and stored as 10mM in 100% dimethyl sulfoxide (DMSO) stock solutions at −20°C. The final concentration of compounds was maintained constantly at 10uM in the experiments, unless mentioned. The mouse anti-Flag (M2, Sigma-Aldrich), and mouse anti-GAPDH (G8140, US Biological, Salem MA) antibodies were used in this study.

### 2.2. Viruses

USA-WA1/2020 and B.1.617.2 strains were obtained from BEI Resources and propagated in Vero E6 cells by infecting the cell monolayer with the virus for 2 h at 37 °C. Unattached virus was removed by washing followed by addition of fresh medium. After 72phi, supernatant was harvested, and the cell debris was removed by centrifugation. The virus was aliquoted and stored at −80 °C until further use. Viral copies in the harvested supernatant were quantified by Reverse Transcriptase qPCR (qRT-PCR) by using standard SARS-CoV-2 genomic RNA with known amounts of SARS-CoV-2 (BEI Resources). SARS-CoV-2 infection assays were performed in BSL-3 containment facility of the University of Nevada, Reno.

### 2.3. Establishment of the Stable cell lines

Permissive HEK293 cells were transfected with RdRp-luciferase reporter and pcDNA3.1. HA or flag-tagged nsp5,7,8,12 constructs (synthesized from Genscript. Inc.) in a 12-well plate, using metafectene and lipofectamine reagent, respectively, as per manufacturer’s instructions. After 24h, cells were selected using 250ug/mL G418 (for 293/RdRp cell lines) or 250ug/mL G418 and 25ug/mL hygromycin (for 293/RdRp-nsp5,7,8,12-Flag cell lines). After 3 weeks of selection, expanded clones were examined for the expression of Flag-tagged nsp12 (RdRp) by western blot using anti-Flag antibody. Same blot was also stained with anti-GAPDH to serve as the loading control. *Firefly* and *Renilla* luciferase reporter gene expression in the cells were measured with a Dual Luciferase Assay System (Promega Corporation, Madison, WI, United States) according to the manufacturer’s instructions. Briefly, 150,000 cells per well were plated in a 12-well plate. For compound treatment, cells were treated with selective inhibitors at a final conc. of 10 μM or DMSO alone (control) for 48h. Cells were then lysed in 100uL of 1X passive lysis buffer and incubated at RT for 15 min to ensure complete lysis. The lysate was transferred to a 96-well microplate, followed by addition of 100μl of the *firefly* luciferase reagent (LARII) with a 10-s equilibration time and measurement of luminescence. Next, 100μl of the *Renilla* luciferase reagent (Stop & Glo) for *firefly* luminescence quenching was added with a 10-s equilibration time and measurement of luminescence. The relative activity of RdRp polymerase was determined by normalizing the level of of *firefly* luciferase activity (FLuc) against that of *renilla* luciferase activity (RLuc), FLuc/RLuc.

### 2.4. Immunoprecipitation and Western blotting

For immunoprecipitation, nearly 5 × 10^6^ cells were harvested and washed with cold 1X PBS, lysed in RIPA lysis buffer (1% NP-40, 50mM Tris [pH 7.5], 1 mM EDTA [pH 8.0], and 150mM NaCl) supplemented with protease inhibitors (1mM phenylmethylsulfonyl fluoride, 1μg/ml aprotinin, 1μg/ml pepstatin, 1μg/mL sodium fluoride, and 1 μg/ml leupeptin), and incubated on ice for 30 min. Cell lysates were sonicated with a probe sonicator and cell debris was removed by centrifugation (12,000 rpm, 10 min at 4°C). The lysates were pre-cleared with protein A+G conjugated sepharose beads (30 min at 4°C). Approx. 5% of the lysate was saved as input sample and the remaining lysate was incubated (while rotating) with anti-flag antibody overnight at 4°C. Immune complexes were captured by incubating with 35μL of protein A+G sepharose beads for 2h at 4°C. The beads bound immune complexes (IP samples) were collected by centrifugation (2,000 rpm, 2 min at 4°C), followed by washing with ice-cold RIPA buffer (2X). The input samples and the IP samples were resuspended in 45μL of SDS-PAGE loading buffer and denatured at 95°C for 5 min, resolved by SDS-PAGE and western blotted onto a 0.45 μM nitrocellulose membrane using standard protocols (Bio-Rad Laboratories, Hercules, CA, United States). Proteins of interest were detected by incubating the membrane with specific antibodies, followed by incubation with appropriate infrared-dye-tagged secondary antibodies (IR-Dye680/800) and scanning with an Odyssey infrared scanner (Li-COR Biosciences, Lincoln, NE, United States).

### 2.5. RNA extraction and RT-qPCR

Relative expressions of FLuc and RLuc gene transcripts in cells were measured by real-time reverse transcription PCR (RT-qPCR). Total mRNA from the cells was extracted using Trizol reagent (Thermo Fisher Scientific, Waltham, MA, United States) according to the manufacturer’s recommendation. The cDNA was made using a high-capacity RNA-to-cDNA Kit (Thermo Fisher Scientific) and amplified on Bio-Rad CFX Connect RT-qPCR detection system (Bio-Rad). Each amplification reaction was performed in 20μL volume containing 10μL of SYBR Green Universal master mix (Bio-Rad), 2.5μL each of forward and reverse primers (2μM) targeting corresponding genes, and 5μL of cDNA or water. Primers used in this study are: **FLuc-F** (5’-ATC GAA GGA CTC TGG CAC AA-3’), **FLuc-R** (5’-CCT ACC GTG GTG TTC GTT TC-3’), **RLuc-F** (5’-AGT GGT GGG CCA GAT GTA AA-3’), and **RLuc-R** (5’-CGC GCT ACT GGC TCA ATA TG-3’). Specificity of amplification was assessed by analyzing the melting curve. The relative mRNA expression level of each target gene (FLuc and RLuc) was normalized to the corresponding beta-actin expression level, and the fold change was calculated using the *ΔΔC_T_* method with respect to cells expressing RdRp reporter plasmid.

For relative viral genome quantification post infection and compound treatment, control or USA-WA1/2020 virus was added onto the Vero E6 cells plated in a 24-well plate (100,000 cells per well) for 2h (37 °C and 5% CO_2_). Post infection, the virus containing medium was replaced with 1 mL fresh medium containing either DMSO (control) or compounds (10μM). The cells were treated with compounds for 72h at 37 °C and 5% CO2. For the detection of viral genomic RNA through qRT-PCR, supernatant from control or compound-treated Vero E6 cells (for intact viral genomic RNA) or from treated cells (for infectious viral genomic RNA) were subjected for total RNA extraction using Trizol reagent (Invitrogen, Carlsbad, CA, USA) via direct-zol RNA extraction kit (Zymo Research), according to the manufacturer’s recommendation. An aliquot of extracted total RNA (5 μL) was used for the relative quantification of viral genomic copies using TaqPath and N1 primer-probe set in a qRT-PCR assay (Thermo Fisher Scientific, Waltham, MA, USA).

### 2.6. Cell Toxicity Assay or Methyl Thiazolyl Tetrazolium (MTT) assay

Cell toxicity due to the compound treatment was determined using the standard colorimetric MTT [3-(4,5-dimethylthiazol-2-yl)-2,5-diphenyltetrazolium bromide] assay (Thermo Fisher Scientific) according to the manufacturer’s instructions. Briefly, 5,000 cells resuspended in the complete DMEM medium were seeded in a costar 96-well microplate and grown overnight. Next day, cells were washed with 1X PBS and treated by adding 100μL of MTT solution (0.5mg/mL) to each well, followed by incubation at 37°C for 4h. The culture medium (100μL) was used as a negative control (blank). After 4h, when intracellular purple formazan crystals were visible under microscope, MTT solution was removed and replaced with 50μL of solubilizing DMSO solution. Cells were incubated at RT for 30 mins in order to dissolve the purple formazan crystals. The absorbance was measured on a microplate reader at a wavelength of 570nm. The background absorbance produced by wells containing medium only (Ab_Sbiank_) was subtracted from all wells and the percent viable cells were calculated using the equation: %viable cells=[(Ab_Ssampie_-Ab_Sbiank_)/ (Ab_Scontrol_-Ab_Sblank_)] *100.

### 2.7. Dose-Response assays

Vero E6 cells were seeded at 100,000 cells per well in a 24-well microplate and allowed to attach for 2h. The cells were infected with USA-WA1/2020 or B.1.617.2 variant for 2h, followed by treatment with dasabuvir at concentrations 100, 33, 11, 3.7, 1.23, and 0.41 μM, in duplicate. After 48 h of incubation at 37 °C with 5% CO2, the supernatant was collected and a fraction (100μl) was used for RNA extraction and rt-qPCR (reverse transcriptase-PCR) to quantify the amounts of SARS-CoV-2 in the supernatant. Another fraction of the supernatant was used for the detection of infectious SARS-CoV-2 through the enumeration of plaque forming units described below. The absolute half-maximal inhibitory concentrations (IC_50_) of dasabuvir for SARS-CoV-2 USA-WA1/2020 and B.1.617.2 were calculated, based on the amounts of residual virus in the supernatant at each concentration, using the Graphpad Prism™.

### 2.8. Plaque Assay

Vero E6 cells (100,000 cells/well) in 1mL medium were seeded in a 24-well culture plate for 2h (37 °C and 5% CO_2_). Next, 20μL and 2μL (10-fold dilution) of virus containing supernatant (post compound treatment) were added onto the cells and the cell plate was incubated for 2h (37°C with 5% CO_2_), followed by the replacement of medium with 1.5% CMC overlay medium. After 7 days of incubation, overlay medium was replaced with 4% paraformaldehyde solution (500μL) and the cells were incubated at room temperature for 30 mins. Fixed cells were stained with 0.2% crystal violet solution in 20% ethanol (500μL) at room temperature for 30 min with gentle rocking after every 10 min. Staining solution was removed, and cells were washed with sterile distilled water (thrice). Plate was inverted and left for drying on absorbent pad for 1-2h. Plaques appeared as clear circles in the purple monolayer of cells (used negative control with no clear circles as reference). For each well, the number of plaques for that dilution were counted and plaque forming unit (PFU per mL) were calculated using the equation-

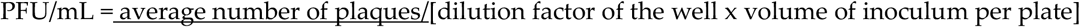

### 2.9. Calculation of Z-Factor and Z’-Factor

In order to evaluate the quality and reproducibility of the screening assay, screening window coefficients, Z- and Z’-factor values were calculated as described previously. Briefly, 50,000 cells per well were plated in a 24-well plate for each group as follows: (1) Control group = cells expressing RdRp reporter plasmid and pcDNA3.1 (control vector), n = 24 wells, treated with 0.025% DMSO; (2) Positive group = cells expressing RdRp reporter plasmid and flag-tagged nsp5, 7, 8, 12 plasmid, n=24 wells, treated with 0.025% DMSO; and (3) Inhibitor group = cells with RdRp reporter plasmid and flag-tagged nsp5, 7, 8, 12 plasmid, n=24 wells, treated with 10uM remdesivir. Post 48h, cells were lysed in 100uL of 1X passive lysis buffer and subjected to dual luciferase reporter assay as mentioned in Section 2.3. The relative activity of RdRp was determined by normalizing the level of of FLuc to that of RLuc (FLuc/RLuc). Z-factor was calculated using the equation: Z-factor = 1 – [(3SD_Control_ + 3SD_Positive_)/| mean_Control_ – mean_Positive_ |], and Z’-factor was calculated using the equation: Z’-factor = 1 – [(3SD_Inhibitor_ + 3SD_Positive_)/| mean_Inhibitor_ – mean_Positive_ |], where SD (the standard deviation) and mean values correspond to the relative FLuc activity obtained from each group.

### 2.10. Statistical analyses

All the experiments were performed three times, and the data is presented as the mean of a triplicate. The error bars represent the standard deviation across three independent experiments. Statistical analyses were performed using Prism 8.0 software (GraphPad Inc.). The p-values were calculated using two-way analysis of variance (ANOVA) tests, and the *p*-value cut offs for statistical significance were *, *p* < 0.1; and **, *p* < 0.01.

## 3. Results

### 3.1. Development of Cell-based SARS-CoV-2 RdRp Reporter Assay

In order to measure the intracellular SARS-CoV-2 RdRp (nsp12) enzymatic activity, we developed a cell-based luciferase reporter system, using similar principle as reported earlier for cell-based HCV RdRp and MERS-CoV RdRp activity assays (17, 18). The schematic of the bicistronic SARS-CoV-2 RdRp reporter construct used in the system, with firefly luciferase (FLuc) and renilla luciferase (RLuc) genes in reverse orientation, designated as p(+)RLuc-(-)UTR-FLuc is shown in Figure 1A. The (–)FLuc is flanked by the 5 - UTR and 3’-UTR of SARS-CoV-2 and ribozyme self-cleavage sequence of the hepatitis delta virus (HDV). The transcription of full length (+)RLuc-(–)UTR-FLuc RNA is catalyzed by the host RNA polymerase Pol II. The generated bicistronic RNA transcripts are further self-cleaved by ribozyme, leading to generation of FLuc RNA in a negative-sense orientation, that is, then transcribed to the positive-sense FLuc RNA by RdRp polymerase for protein synthesis. Thus, intensity of the measured FLuc signal is proportional to the level of intracellular RdRp polymerase activity. In addition, in the presence of RdRp activity, only the amounts of synthesized (+) Fluc RNA gets amplified whereas the levels of RLuc RNA (serving as internal control of transcription and translation) remains unaffected. To transiently express SARS-CoV-2 RdRp in mammalian cells, we constructed flag-tagged nsp5, 7, 8,12 vector as shown in Figure 1B. The construct expressed RdRp as a polypeptide with other accessory proteins, namely nsp7, nsp8 (required for efficient RdRp activity) and 3CL^pro^ (nsp5, with autoproteolytic activity). A flag tag introduced at the 3’end of nsp12 (RdRp) helped in determining the expression and cleavage from the polypeptide.

**Figure 1.**
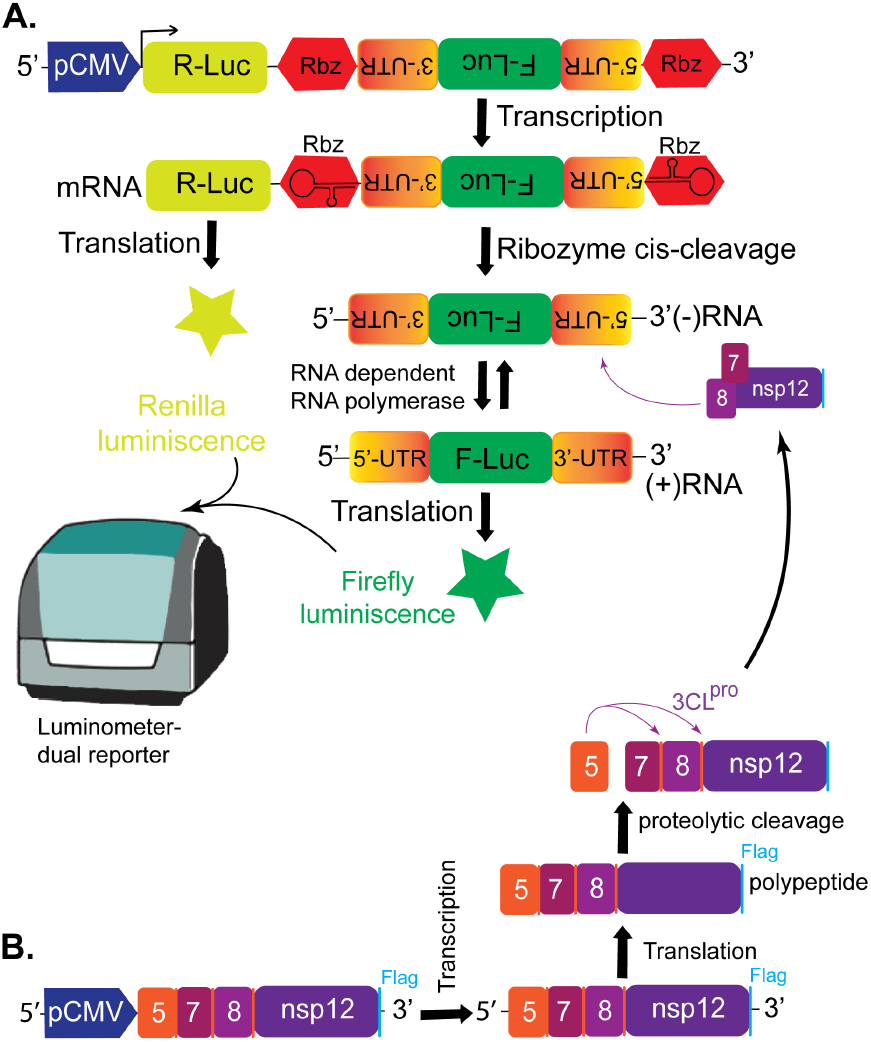
Cell-based SARS-CoV-2 RdRp Activity Reporter. A) Schematic diagram of the bicistronic SARS-CoV-2 RdRp reporter construct p(+)RLuc-(-)UTR-FLuc.The construct contains *renilla* luciferase (RLuc) gene cloned in positive direction under CMV promoter, and the *firefly* luciferase (FLuc) gene under the control of 5’-untranslated region (UTR), the binding site of RdRp for replication (positive sense RNA). FLuc gene is flanked by 5’-UTR and 3-UTR of SARS-CoV-2 and the hepatitis delta virus (HDV) ribozyme self-cleavage sequence. RLuc serves as an internal control to normalize FLuc activity. Expression of RdRp in trans will control FLuc levels and compounds/mutations altering RdRp activity will be reflected by reduction in FLuc levels. (B) Schematic representation of the plasmid construct expressing SARS-CoV-2 RdRp (nsp12), accessory proteins (nsp7, and nsp8) and viral 3CL^pro^ protease (nsp5), required for proper cleavage of the proteins. Cleavage of the proteins is confirmed by the expression of flag-tagged nsp12 (RdRp) in an immunoblotting assay.

### 3.2 Determination of SARS-CoV-2 RdRp polymerase activity

To test our SARS-CoV-2 RdRp reporter system and the expression and the activity intracellular SARS-CoV-2 RdRp, we co-transfected RdRp-reporter construct with pcDNA3.1 (control) or varying amounts (1μg and 3μg) of flag-tagged nsp5,7,8,12 expressing plasmid in HEK293 cells. At 48h post-transfection, we tested the expression of flag-tagged SARS-CoV-2 RdRp in the cells by western blotting with anti-flag antibody, along with anti-GAPDH for the loading control (Figure 2A). A dose-dependent increase in the RdRp protein expression was observed in the cells as confirmed by bands at 110kDa. The absence of corresponding band in the cells transfected with empty vector confirmed the specificity of Rdp expression. The dose-dependent increase in the FLuc and FLuc/RLuc activities is expected to be proportional to the increase in levels of (+) FLuc RNA transcripts, which is dependent on RdRp polymerase activity. In order to prove this hypothesis, we performed dual luciferase reporter assay and rt-qPCR on HEK293 cells co-transfected with the above combination of plasmids. As seen in Figure 2B, both the FLuc and the relative FLuc (Index value) activities were increased by the expression of flag-tagged SARS-CoV-2 RdRp in a dose-dependent manner, while the increase in RLuc activities was comparable. The rt-qPCR using primers specific to the FLuc and RLuc genes, with beta-actin serving as the internal control, showed an increase in the FLuc mRNA expression level. In addition, increase in the relative FLuc/RLuc mRNA expression levels was directly proportional to increase in amounts of flag-tagged RdRp present in HEK293 cells (Figure 2C). No significant increase in the levels of RLuc RNA was observed for the transfected cells, conforming that RdRp did not affect the levels of RLuc, as expected.

**Figure 2.**
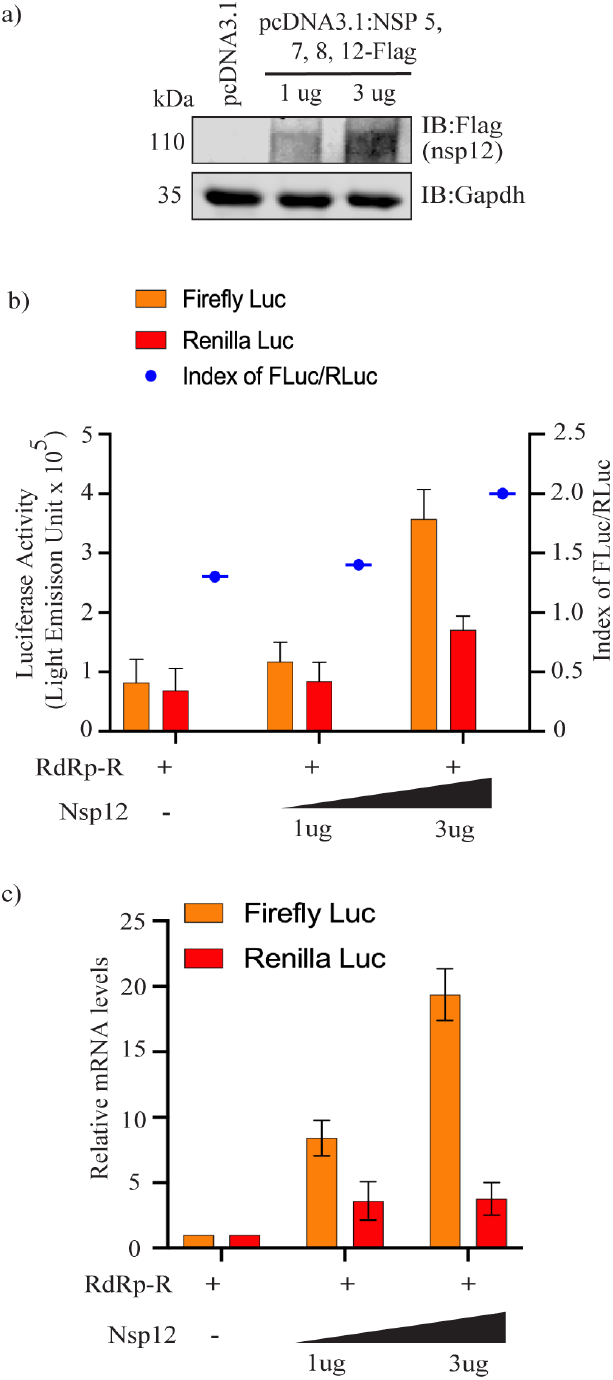
Transient reporter assay of SARS-CoV-2 RdRp polymerase. (A) HEK293L cells expressing SARS-CoV-2 RdRp reporter with either control pcDNA3.1 or flag-tagged nsp5,7,8,12 constuct were subjected to luminescence detection with the dual luciferase reporter assay kit (Promega) 48hpt, and the FLuc, RLuc and ratio of FLuc/RLuc (Index) activities were determined. (B) Transiently transfected cells were also subjected to immunoblotting with anti-flag antibody 48h posttransfection. (C) Total RNA was extracted and FLuc mRNA and RLuc mRNA expression levels were assayed by RT-qPCR.

### 3.3. Generation and Characterization of HEK293 cells stably expressing SARS-CoV-2 RdRp reporter construct

Since the FLuc levels, indirectly measuring the RdRp activity, was significantly enhanced in the transient assay described above, we generated HEK293 cells stably expressing SARS-CoV-2 RdRp reporter and pcDNA3.1 (control) or flag-tagged nsp5, 7, 8, and 12 polypeptide by co-transfecting RdRp-reporter with either pcDNA3.1 or flag-tagged nsp5, 7, 8, 12 construct. The stable expression of RdRp polymerase aids in development of reporter system for a high-throughput screening (HTS) platform. Transfected cells were selected with 250ug/mL G418 (293-RdRp cells) or 250ug/mL G418 and 25ug/mL hygromycin (293-RdRp-nsp5,7,8,12-flag cells) at 24h post-transfection. After 3-weeks of selection, cells were analyzed for luminometric activity using the dual luciferase reporter assay kit (Figure 3A), and RdRp protein expression by immunoblotting assay using anti-flag antibody (Figure 3B). For the reporter assay, equal number of 293-RdRp and 293-RdRp-nsp5, 7, 8,12-flag cells were treated with DMSO for 48h, and assayed with dual luciferase reporter assay. In addition, we treated equal number of 293-RdRp-nsp5, 7, 8,12-flag cells with remdesivir, a known SARS-CoV-2 RdRp inhibitor, to further validate our SARS-CoV-2 RdRp reporter system. As seen in Figure 3A, the calculated FLuc/RLuc ratio (Index value) in the cells constitutively expressing RdRp reporter and the polypeptide, was higher (Fig. 3A, RdRp Nsp-DMSO bars) than cells harboring RdRp reporter without the nsp5, 7, 8,12-flag (Fig. 3A, RdRp-DMSO bars). Importantly, addition of remdesivir reduced the levels of FLuc or relative FLuc/RLuc ratios (index value), with respect to the DMSO-treated set (Fig. 3A, RdRp Nsp-Rem bars), as expected. Western blot analysis of these cells confirmed the expression of flag-tagged SARS-CoV-2 RdRp protein, indicated by the appearance of expected size band at 110kDa (Figure 3B) in 293-RdRp-nsp5,7,8,12-flag cells (mock-as well as resdesivir treated sets). 293-RdRp cells lacked the detection of RdRp, as expected. Relative quantification of the bicistronic genes (FLuc and RLuc) copy number from stable cells was performed using quantitative rt-PCR with primers corresponding to the FLuc gene and RLuc, with levels of the cellular β-actin gene as the normalization control. The increase in FLuc levels in the 293 cells, constitutively expressing SARS-CoV-2 RdRp reporter and flag-tagged RdRp protein was proportional to the increase in the mRNA levels of (+) FLuc (generated by RdRp), as compared to RdRp reporter cells lacking RdRp expression (Figure 3C, lane 1 and 2). The levels of (+) FLuc RNAs were also quantified after treatment with remdesivir. As seen in Figure 3C, treatment with RdRp inhibitor significantly reduced the mRNA levels of (+) FLuc. This confirmed that we have developed a cell-based reporter system to quantitatively assess the intracellular SARS-CoV-2 RdRp activity.

**Figure 3.**
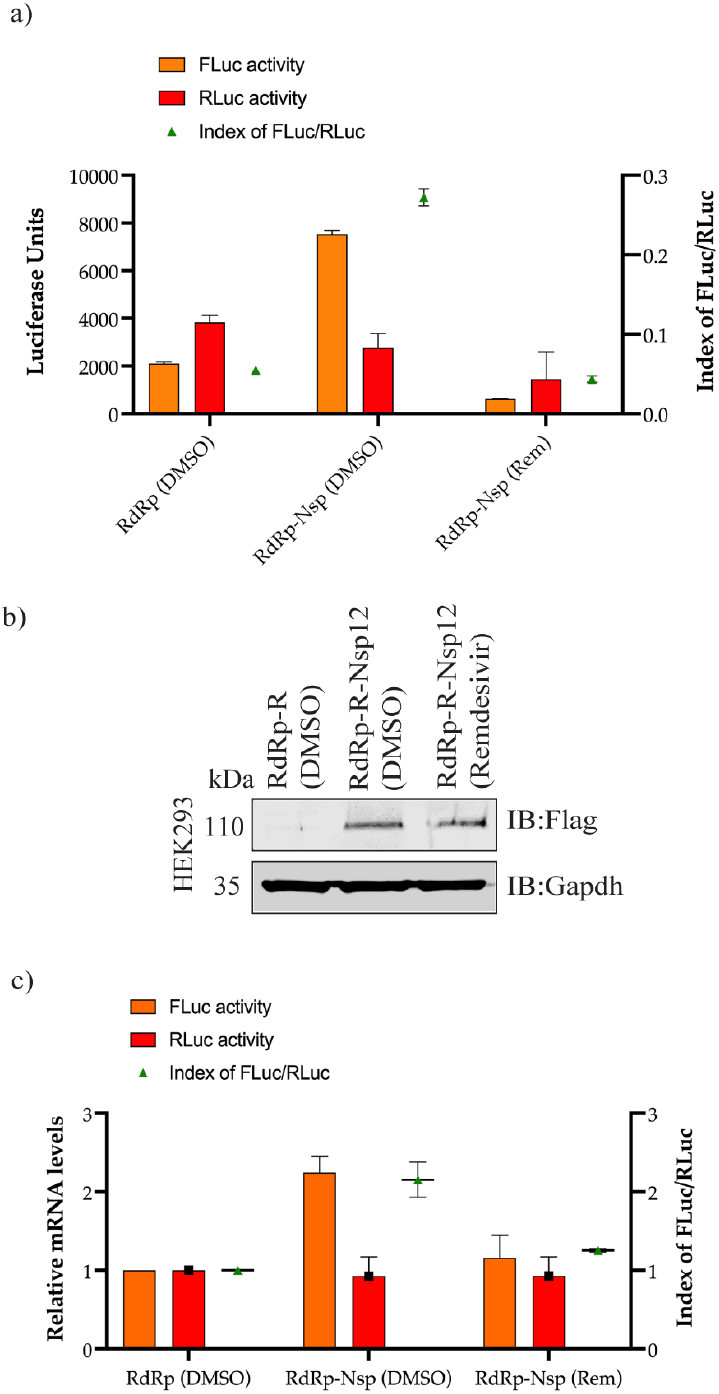
Characterization of 293 cells stably expressing SARS-CoV-2 RdRp reporter. Cells stably expressing the SARS-CoV-2 RdRp reporter and pcDNA3.1 or flag-tagged nsp5,7,8, and 12 constuct were mock (DMSO) treated or treated with remdesivir (a potent anti-RdRp inhibitor), followed by luminescence detection with the dual luciferase reporter assay, and the relative FLuc/RLuc ratio (Index) was detected after 48h. (B) Cells were also subjected to immunoblotting with anti-flag antibody 48h post-transfection, and C) Total RNA was extracted and relative FLuc/RLuc RNA expression levels were quantified by RT-qPCR, using luciferase gene specific primers and beta-actin, as an internal control.

### 3.4. Evaluation of Reporter Assay System for High-Throughput Screening (HTS) of SARS-CoV-2 RdRp Inhibitors

To determine if the cell-based reporter assay is suitable for HTS applications, we calculated screening window coefficients, Z-, and Z’-factors using Zhang’s formulae (19). The Z-, and Z’-factors are HEK293L-RdRp-R cells widely used parameters to evaluate and validate the reliability and reproducibility of screening assay systems as HTS platforms. To calculate the screening coefficients, HEK293 cells stably expressing SARS-CoV-2 RdRp reporter system and pcDNA3.1 vector (control group) or flag-tagged nsp5, 7, 8, and 12 vector (positive group) were treated with DMSO (0.025%) for 24h. HEK293 cells stably expressing SARS-CoV-2 RdRp reporter and flag-tagged nsp5, 7, 8, and 12 vector were used for determining the luciferase levels without and with remdesivir (10μM, inhibitor group) treatments for 24h thrugh dual luciferase reporter assay. The Z-factor (specificity of the assay for SARS-CoV-2 RdRp activity) and Z’-factor (applicability of remdesivir as a positive control), were calculated using the relative FLuc activity obtained from control and positive, and positive and inhibitor groups, respectively. We obtained a Z-factor and Z’-factor values of 0.732 and 0.789 respectively, indicating the HEK293 cell-based reporter assay is excellent for assaying RdRp activity of SARS-CoV-2, and confirming the use of remdesivir as a positive control for SARS-CoV-2 RdRp specific inhibitors in HTS assays (Figure 4). Thus, the target-specific HEK293 cell-based assay system targeting the RdRp of SARS-CoV-2 shall provide excellent platform to facilitate identification of efficacious SARS-CoV-2 RdRp antivirals.

**Figure 4.**
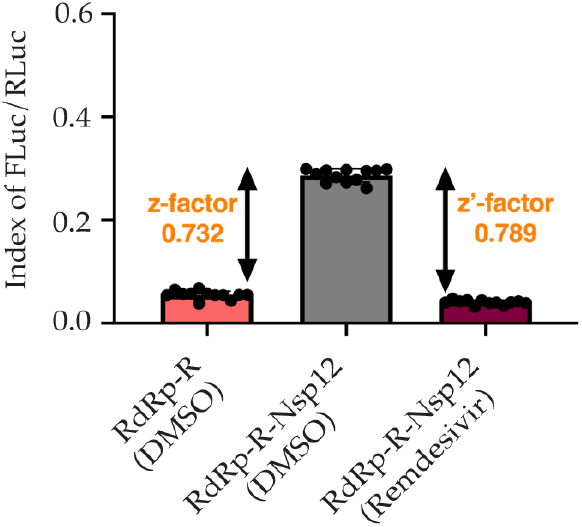
Evaluation of HEK293 cell-based SARS-CoV-2 RdRp Reporter System for HTS assays. The experimental groups were characterized as: control group (n=24)-cells expressing SARS-CoV-2 RdRp reporter and pcDNA3.1 plasmids treated with DMSO (0.025%); positive group (n=24)-cells expressing SARS-CoV-2 RdRp reporter and flag-tagged nsp5, 7, 8, and 12 plasmids treated with DMSO (0.025%); and inhibitor group (n=24)-cells expressing SARS-CoV-2 RdRp reporter and flag-tagged nsp5, 7, 8, and 12 plasmids treated with remdesivir (10μM). The Z- and Z’-factors were calculated for HEK293 cell-based SARS-CoV-2 RdRp reporter systems, using control and positive, and positive and inhibitor groups in each 24-well plate. According to Zhang’s formula, the Z-and Z’factors for HEK293 cell based reporter assay were calculated as 0.732, and 0.789, respectively, indicating assay system has the required robustness and reproducibility for HTS extracted and FLuc mRNA and RLuc mRNA expression levels were assayed by qRT-PCR.

### 3.5. Screening of RdRp inhibiting antivirals

We further evaluated if the optimized SARS-CoV-2 RdRp reporter system allows for the screening of inhibitors against SARS-CoV-2 RdRp activity and virus replication *in vitro.* To this end, we determined the efficacy of commercially available viral DNA or RNA polymerase inhibitors as potential anti-virals for SARS-CoV-2, including rhoifolin, ribavirin, dasabuvir, penciclovir, and cytidine-5’-triphosphate (C5’T) and DMSO (control) using our HEK293 cell-based SARS-CoV-2 RdRp reporter system. These inhibitors were added to the stable cells at a final conc. of 10μM in medium with 10%FBS. After 48h, cells were lysed and subjected to dual luciferase reporter assay and FLuc/RLuc ratio (index) were determined, with the levels of FLuc activity reflecting the anti-RdRp activities of the compounds. As shown in Figure 5A, a marked reduction in the FLuc/RLuc ratio was observed in cells treated with dasabuvir and C5’T, whereas, rhoifolin- and penciclovir-treated cells only showed a slight reduction in the FLuc/RLuc values or SARS-CoV-2 RdRp activity, with respect to the DMSO-treated cells. Interestingly, cells treated with ribavirin in combination with dasabuvir exhibited higher anti-RdRp activity when compared to cells treated with ribavirin alone. This data confirmed that our reporter system stably expressing SARS-CoV-2 RdRp can be used for selective screening of inhibitors of SARS-CoV-2 replication within the cellular environment. In order to determine whether these antivirals have cytotoxic effects on those treated cells, which may also contribute to alter levels of FLuc/RLuc, we performed cell viability assay using MTT analysis. This colorimetric assay provides a sensitive and accurate method for the determination of cell viability based on the generation of an insoluble formazan (purple color) from water-soluble tetrazolium MTT dye. As shown in Figure 5C, no detectable cytotoxic effects were observed in the MTT analysis, when the cells were treated with 10μM conc. of these selected anti-virals for 48h. Hence, this indicated that our HEK293 cell-based SARS-CoV-2 RdRp reporter system will be instrumental in screening additional novel inhibitors of SARS-CoV-2 RdRp activity.

**Figure 5.**
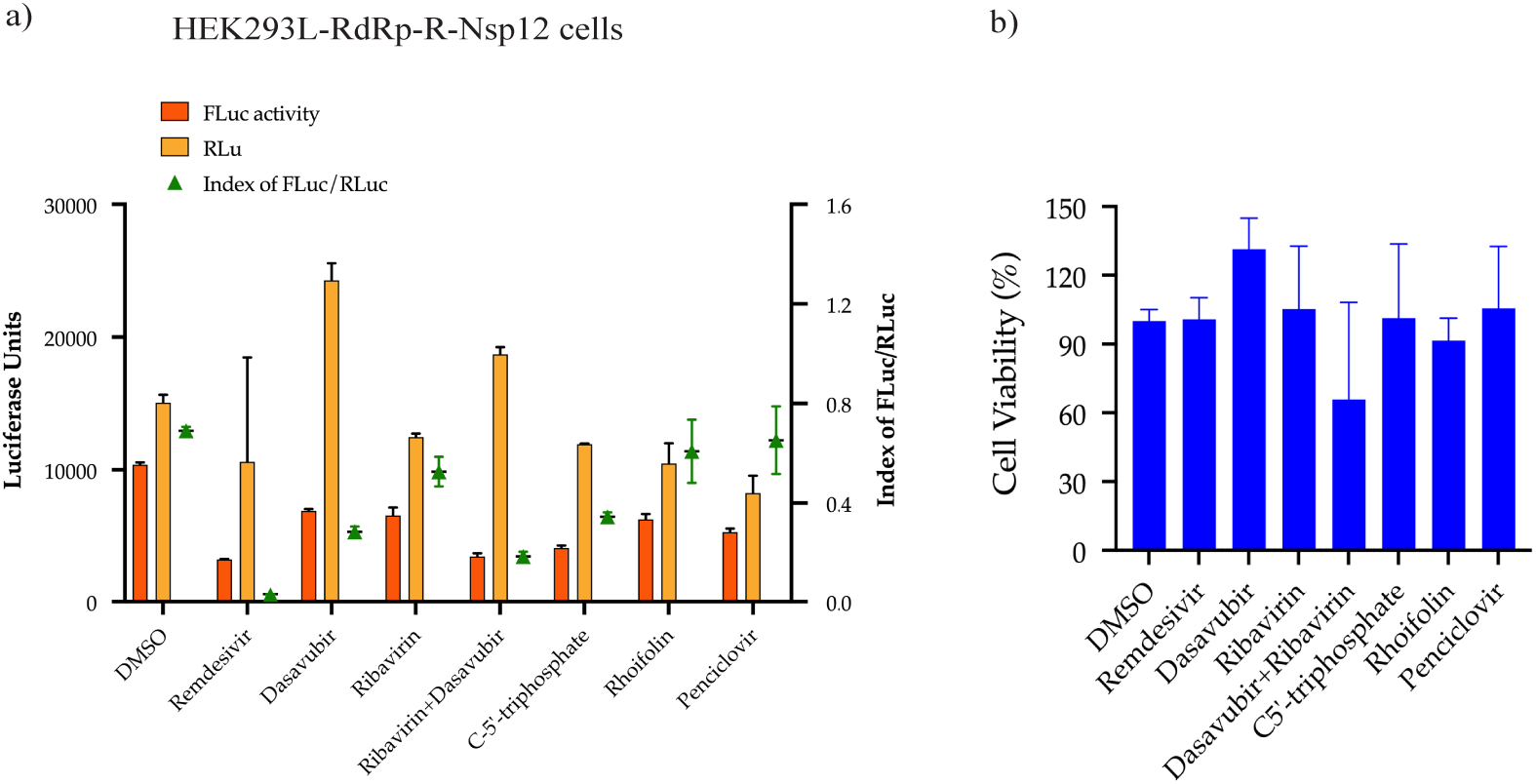
Screening of the SARS-CoV-2 RdRp inhibiting compounds through cell-based reporter system. (a) HEK293L cells with RdRp reporter system were treated with DMSO (control) or 10μM of test compounds and subjected to luminescence detection with the dualluciferase reporter assay kit post 48h treatment. Firefly (FLuc) and Renilla (RLuc) luciferase values are plotted and the ratios of FLuc and RLuc (Index values), indicator of RdRp inhibition are shown. (b) Toxicity of inhibitors at 10μM concentration on HEK293 cells was analyzed by MTT assay, 48h post-treatment. Results are presented as percent reduction with reference to DMSO from three independent experiments performed in triplicates.

### 3.6 Effective Inhibition of SARS-CoV-2 Replication by Dasabuvir in vitro

We further investigated the antiviral activity of chosen compounds against SARS-CoV-2 replication *in vitro.* To this end, USA-WA1/2020 (1.6 × 10^6^ TCID50/mL; Lot # 70036318) and B.1.617.2 (6.5 × 10^5^ TCID50/mL^2^; Lot # 70045238) variants were obtained from BEI Resources and propagated in Vero E6 cells. Viral titers were determined after 3 days by rt-qPCR using N1 primer-probe set. The antiviral activities of these chosen RdRp inhibitory compounds were determined by infecting Vero E6 cells with USA-WA1/2020 virus for 2h (37 °C and 5% CO_2_), and subsequently treating the cells with DMSO (control) or test compounds (10μM). After 3 days of incubation, viral RNA was extracted from the supernatant as well as the cells and genomic SARS-CoV-2 RNA was quantified using rt-qPCR assay (Figure 6). Intracellular genome copies of the virus in the cells treated with different compounds were determined relative to the copies in the DMSO treated cells (Figure 6a). Cell free genome copies of the virus in the supernatants were determined by using a standard curve generated with a known amounts of SARS-CoV-2 genomic RNA (BEI Resources, NR-52285, Lot # 70033700) (Figure 6b). We detected a significant decrease in the number of virus copies in the supernatants of cells treated with dasabuvir and its combination with Ribovarin. Remdesivir, showed a reduction in the viral genome copies in the supernatant, as expected. Next, the supernatants from compound treated cells were evaluated for the presence of viable virus through plaque assay (Figure 6d). Monolayer of Vero E6 cells were inoculated with 20 μl and 2 μl of the collected supernatant for 2h and overlaid with 1.5% CMC containing DMEM medium. Seven days later, cell monolayers were fixed with formaldehyde and stained with crystal violet. Plaques were enumerated by direct visualization and PFUs values were reported as plaque-forming units per mL (PFU/mL) calculated through the Poisson distribution. Both dasabuvir-treated cells and dasabuvir/ribavirin-treated cells completely blocked the SARS-CoV-2 replication and propagation, as evident from absence of plaques (PFU/mL). Quantitative analysis of the antiviral activity of dasabuvir against SARS-CoV-2 replication was determined by calculating the half-maximal inhibitory concentration (IC_50_). Vero E6 cells were infected with USA-WA1/2020 or B.1.617.2 virus (50μL) and treated with varying concentrations of dasabuvir (serial dilutions ranging from 100uM to 0.45 μM), along with untreated cells serving as a control. After 48 h, SARS-CoV-2 RNA levels in the supernatants were quantified through the standard curve of Ct values and viral genome equivalents. The IC_50_ values of dasabuvir for WA1 and B.1.617.2 were determined by plotting the viral genome detected in the supernatants at each concentrations using GraphPad. Dasabuvir treatment reduced the levels SARS-CoV-2 in the supernatants of cells infected with WA1 and B.1.617.2 viruses (calculated IC_50_ values of 9.47 μM and 10.48 μM, for WA1 and B.1.617.2, respectively), indicating potential anti-SARS-CoV-2 activity of dasabuvir (Figure 6e and 6f). Taken together, we demonstrated that dasabuvir exhibits antiviral activity against SARS-CoV-2 and reduced SARS-CoV-2 replication efficiently in an in-vitro cell culture model.

**Figure 6.**
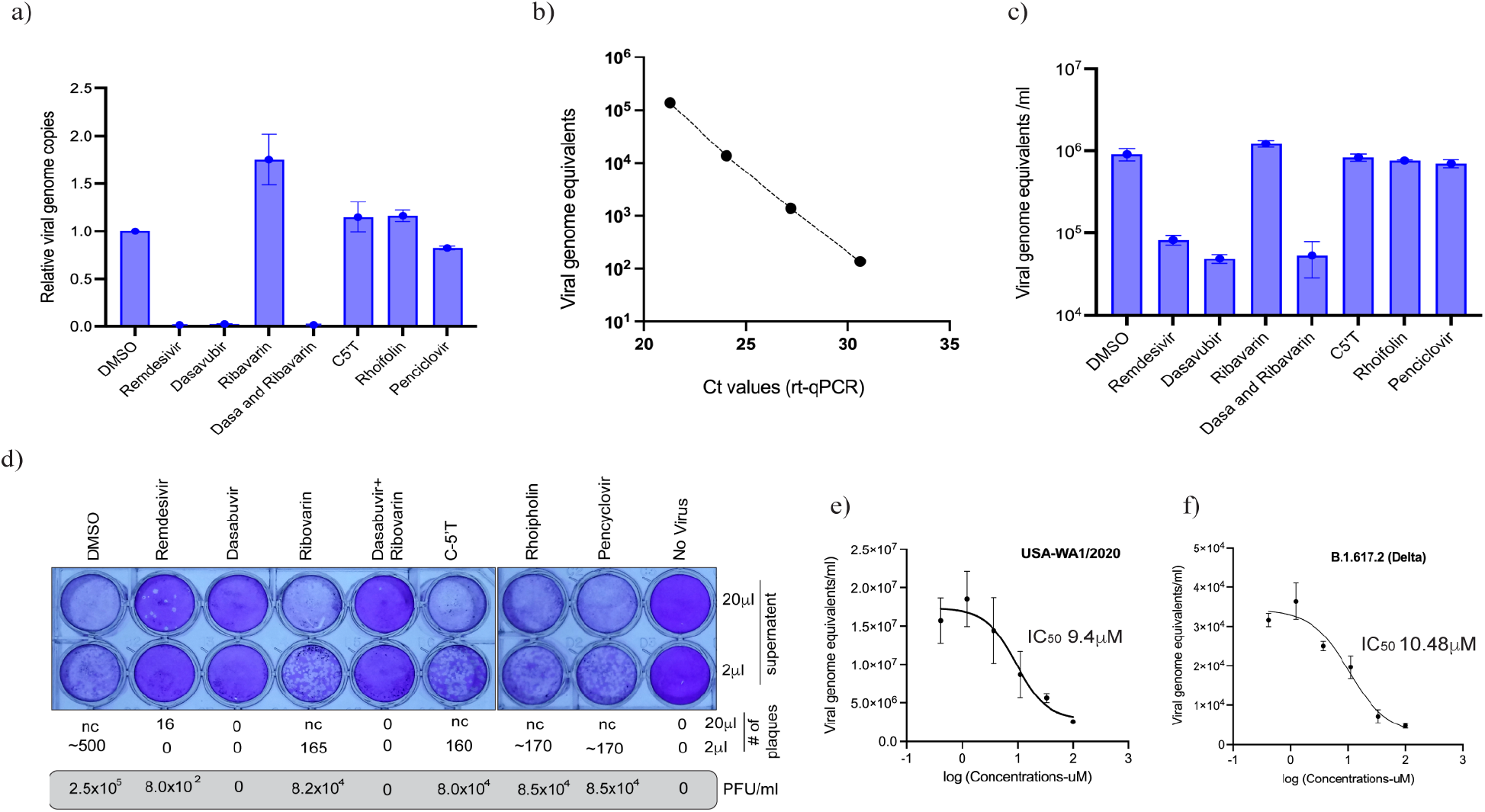
Dasabuvir inhibits SARS-CoV-2 replication in Vero E6 cells. (a) Vero E6 cells were infected with USA-WA1/2020 and subsequently treated with DMSO (control) or 10 μM of the test compounds for 48h. a) Extracted RNA from the cells were used for the quantitation of intracellular SARS-CoV-2 genome through rt-PCR and plotted after normalization with housekeeping gene. (b) Standard curve of known viral genome copies and Ct values for calculating the genome equivalents of SARS-CoV-2 in the supernatants. (c) Quantitation of cell free SARS-CoV-2 genomes in the supernatants of compound treated cells through rt-PCR on extracted RNA. Genome copies per ml was calculated based on the Ct values using the standard curve. (d) Determination of infectious virus in the supernatant of treated cells through plaque assay. 20 μl and 2 μl of the supernatants from mock (no virus) and USA-WA1/2020 virus infected cells and treated with different compounds were used for the plaque assay. Plaques were counted for determining the per ml plaque forming units (PFU). nc= not countable. IC50 value of dasabuvir against SARS-CoV-2 was determined by measuring a reduction in the viral genome copies in the supernatant of (e) USA-WA1/2020 and (f) B.1.617.2 (Delta) infected cells following treatment with the varying amounts of dasabuvir.

## 4. Discussions

Emergence of highly pathogenic HCoVs, especially, SARS-CoV, MERS-CoV, and recently SARS-CoV-2, have threatened public health security of almost all countries in the world. Although COVID-19 prophylactic vaccines currently authorized for use in United States are effective against COVID-19 infection, the risk of SARS-CoV-2 infection in a fully vaccinated person can’t be eliminated. Besides, new variants of SARS-CoV-2 circulating around the globe have been reported to evade antibodies that neutralize the original SARS-CoV-2 variant, making them less effective as the new variants become dominant. Hence, an urgent need remains to rapidly develop, and validate new and existing efficacious anti-SARS-CoV-2 drugs to limit the mortality and morbidity of severe COVID-19 infections.

Among other crucial viral proteins, SARS-CoV-2 encoded RdRp plays essential roles in virus life cycle and is designated as a potential target for anti-SARS-CoV-2 drug development. Owing to identical sequence among HCoVs and lack of human homolog, SARS-CoV-2 RdRp allows for specific and selective RdRp inhibition without affecting other cellular protein. To exploit SARS-CoV-2 RdRp as a potential drug target, it is essential to have a stable, cell-based, target-specific functional assay that allows quantitative measurement of RdRp polymerase activity. Such an assay will facilitate determination of potential RdRp-specific inhibitors. Notably, a cell-based assay stably expressing RdRp protein would further help eliminate both cytotoxic and membrane impermeable inhibitors. Despite the importance of a cell-based RdRp activity assay, an optimal cell-based assay stably expressing RdRp protein is currently unavailable for assaying the polymerase activity of SARS-CoV-2 RdRp.

While this manuscript was under preparation, Zhao *et al* reported a cell-based CoV-RdRp-Gluc assay for the identification of SARS-CoV-2 RdRp inhibitors (20). Culture supernatants of HEK293T cells transiently transfected with CoV-Gluc reporter along with nsp12, nsp7, nsp8, nsp10, and nsp14 expressing plasmids were subjected to Gluc activity assay. Although the assay successfully evaluated the efficacy of seven nucleotide analog compounds, however, the lack of steady RdRp expression adds variability in the gene expression and limits the immediate utilization of the developed assay in HTS applications. Here, we report the development of a simple and efficient HEK293 cell-based reporter assay system with steady RdRp protein expression for measuring the SARS-CoV-2 RdRp activity *in cells*. The system consists of the bicistronic SARS-CoV-2 RdRp reporter construct, p (+) RLuc-(-) UTR-FLuc and flag-tagged nsp5, 7, 8, and 12 plasmids. The FLuc gene expression represents the intracellular RdRp activity, whereas the RLuc gene expression serves as an internal control. This system allows a rapid and quantitative measurement of SARS-CoV-2 RdRp activity in a cell-based format.

We quantified the accuracy of the system by screening known viral anti-polymerase drugs such as rhoifolin, penciclovir, ribavirin, dasabuvir, and cytidine-5’-triphosphate, as effective inhibitors of SARS-CoV-2 RdRp activity. Among them, rhoifolin, a flavonoid earlier shown to inhibit SARS-CoV 3CL^Pro^ (21) and penciclovir, a guanosine analogue that is predicted to bind to SARS-CoV-2 RdRp via molecular docking studies (22), did not significantly inhibit SARS-CoV-2 RdRp activity. Ribavirin, a broad-spectrum antiviral prevents viral RNA synthesis by depleting cellular guanosine triphosphate (22). The incorporation of ribavirin triphosphate by RdRp has been shown to result in lethal viral mutagenesis (23, 24). Our results illustrated that ribavirin only partially inhibited SARS-CoV-2 RdRp activity. Dasabuvir, a non-nucleoside HCV NS5B inhibitor and a derivative of benzothiadiazine (25) suppressed SARS-CoV-2 RdRp activity at the conc. of 10 μM. Interestingly, ribavirin when used along with dasabuvir at 10 μM conc, did not alter the inhibitory effect of dasabuvir on SARS-CoV-2 RdRp activity. Taken together, we have developed a convenient, accurate, reproducible, and HTS-compatible reporter-based assay for rapid detection of inhibitors of anti-SARS CoV-2 RdRp activity without the need of culturing infectious SARS-CoV-2 virus. The described reporter assay system would be instrumental in screening and validating the in silico and virtually designed inhibitors and help expedite identification of novel anti-COVID-19 therapeutic agents in the future.

## Acknowledgments

SARS-CoV-2 viruses USA-WA1/2020 (NR-52281) and B.1.617.2 (NR-55611) were obtained from BEI Resources, NIAID, NIH.

